# Potential combined pro-cognitive, anxiolytic and antidepressant properties of novel GABAA receptor positive modulators with preferential efficacy at the α5-subunit

**DOI:** 10.1101/332908

**Authors:** Thomas D. Prevot, Guanguan Li, Aleksandra Vidojevic, Keith A. Misquitta, Corey Fee, Anja Santrac, Daniel E. Knutson, Michael R. Stephen, Revathi Kodali, Nicolas M. Zahn, Leggy A. Arnold, Petra Scholze, Janet L. Fisher, Bojan D. Marković, Mounira Banasr, Jim Cook, Miroslav Savic, Etienne Sibille

## Abstract

Altered γ-aminobutyric acid (**GABA**) function is consistently reported in psychiatric disorders, normal aging and neurodegenerative disorders, and reduced function of somatostatin - expressing GABA interneurons is associated with both mood and cognitive symptoms. Somatostatin-neurons signal in part through α5-subunit containing GABA_A_ receptors (α5-GABAA-Rs) which are localized in brain regions implicated in emotion and cognition. We hypothesize that enhancing α5-GABAA-R activity has therapeutic potential for both mood and cognitive symptoms in stress-based and aging rodent models.

We synthesized four novel imidazobenzodiazepine (IBZD) amide ligands, tested them for positive allosteric modulation at α5-GABAA-R (α5-PAM), pharmacokinetic properties, and for anxiolytic and antidepressant activities in adult mice. Pro-cognitive activity was tested in adult mice submitted to chronic stress and in old mice. Diazepam (DZP), with broad PAM activity at GABAA-Rs, was used as a control.

Three novel IBZD amide ligands (GL-II-73, GL-II-74 and GL-II-75) demonstrated adequate brain penetration, affinity and α5-PAM activity, and metabolic stability for in vivo studies. GL-II-73/74/75 showed significant anxiolytic and antidepressant efficacies in adult mice. GL-II-73 and GL-II-75 significantly reversed cognitive deficits induced by stress or occurring throughout normal aging. This activity was maintained after sub-chronic administration for GL-II-73. In contrast DZP displayed anxiolytic but no antidepressant or pro-cognitive activities.

We demonstrate for the first time the potential for combined anxiolytic, antidepressant and pro-cognitive therapeutic, mediated by newly designed IBDZ amide ligands with efficacy at α5-GABAA-Rs. These results suggest a novel therapeutic approach targeting both mood and cognitive symptoms in depression and/or aging.

## INTRODUCTION

Altered levels and signaling of gamma-aminobutyric acid (**GABA**), the major inhibitory neurotransmitter, is frequently reported in multiple psychiatric disorders (depression (1,2), schizophrenia (3), anxiety disorder (4)), normal ageing (5,6) and neurodegenerative disorders (Alzheimer’s disease, AD) (6,7) in multiple brain regions including the prefrontal cortex (8), anterior cingulate cortex (9) and amygdala (10). Postmortem studies have associated these GABAergic deficits to a reduction in number and/or loss of function of somatostatin-expressing GABA interneurons in multiple brain regions (see (6) for review). Previous studies showed that impaired inhibition from somatostatin-expressing GABA neurons mediated emotionality (11-13) and cognitive deficits (14-16).

GABA receptors are represented by two classes: GABA_A_ and GABA_B_ receptors. GABA_A_ receptors (**GABAA-Rs**) are composed of multiple subunits (α, β, γ, δ, ε, θ, π and ρ) assembled into pentamers (17) forming a chloride channel. GABA released from somatostatin interneurons are thought to signal, in large part, through the α5-containing GABAA-Rs, due to their preferential localization to extrasynaptic distal dendritic regions regulating tonic inhibition (18-21), and additionally through the ubiquitously-expressed α1-GABAA-Rs. Benzodiazepines (**BZs;** such as diazepam, **DZP**) and imidazobenzodiazepines (**IBZDs**, such as flumazenil) represent the well-established classes of therapeutics acting on GABAA-Rs. BZs are prescribed to alleviate the burden of anxiety (22), but have debatable efficacy on the core symptoms of depression such as anhedonia (23). BZs modulate GABAA-R activity as non-selective positive allosteric modulators (**PAMs**), via binding between the γ2 and either the α1, α2, α3 or α5 subunit (24). This broad receptor activity contributes to significant side-effects (sedation, hypnosis, ataxia, addiction, amnesia), limiting their therapeutic potential (25). However, the relations are not straightforward; for instance, high activity at α1-GABAA-Rs induces sedation and contributes to amnesia, but the same receptor subtype is also proposed to be implicated in cognition (26). The BZ anxiolytic properties are mediated predominantly by α2-GABAA-Rs (27). α5-GABAA-Rs have predominant distribution in the neocortex and hippocampus (28) suggesting a role in cognition and emotion (29,30). As α5-PAMs can alleviate behavioral emotionality in a mouse model of schizophrenia (31,32) and depression (33), or exert pro-cognitive efficacy in old rats (34), we hypothesize that bypassing the somatostatin/GABA interneurons deficiency through enhanced α5-GABAA-R activity may represent a novel target for combined therapeutic actions on mood and cognition.

Based on an IBZD structure (a BZ-imidazol hybrid) we developed ethyl esters of IBZDs with a C(4) R-CH3 substituent that bind preferentially to the α5β2/3γ2 GABAA-Rs (35-37) and that are active in models targeting schizophrenia (31,32) and depression (33) but quickly metabolized (38). Because of their improved metabolic stability and bioavailability as compared to esters, amides are commonly-used replacements (39). Hence, a series of amides were designed that fit the pharmacophore/receptor model (37), with increased binding affinity and efficacy at α5-GABAA-Rs, compared to corresponding esters (40). We describe here the synthesis, pharmacokinetic, functional and behavioral characteristics of four IBZD amide ligands and we predicted that the novel α5-PAMs will have significant antidepressant potential and pro-cognitive activity in adult and old mice, with limited adverse effects.

## METHODS AND MATERIALS

Detailed methods are provided in the supplements.

### Chemistry

Novel IBZD amide analogs were synthesized based on the structures of the existing ligands relatively selective for α5-GABAA-Rs, i.e. SH-053-2’F-R-CH3 and MP-III-022 (41). These new ligands contain a chiral (R) C(4)-methyl group and a 2’F pendent phenyl ring, as general structural requirements for improved selectivity at the α5-subtype based on the pharmacophore model (37). All four amide analogs, referred herein as GL-II-73, GL-II-74, GL-II-75, and GL-II-76, were prepared from the SH-053-2’F-R-CH3 following steps described in the **Supplementary Methods** and presented in **Supplementary Figure S1**.

### Electrophysiological recordings

Electrophysiological recordings were performed as described in (42) (**Supplementary Methods)**. Briefly, HEK-293T cells were transiently transfected with mammalian clones of GABAA-Rs. Currents in response to GABA and GABA+ modulators (EC_3-5_ GABA) were measured using voltage-clamp recordings in the whole-cell configuration.

### Binding studies

Binding studies were performed on human embryonic kidney (HEK) 293 cells following the methods described in (39) (**Supplementary Methods**). HEK293 cells were transfected with cDNAs encoding rat GABAA-R subunits subcloned into pCI expression vectors. To determine the equilibrium binding constant KD of [3H]-flunitrazepam for the various receptor-subtypes, membranes were incubated with various concentrations of [3H]-flunitrazepam in the absence or presence of 5mM DZP. Drug concentrations resulting in half maximal inhibition of specific [3H]-flunitrazepam binding (IC50) were converted to Ki values by using the Cheng-Prusoff relationship (43).

### Animals

Young (2-3 months) or old (21-22 months) C57BL/6 mice were kept in normal housing conditions with a 12hr light-dark cycle, water and food *ad libitum*. Prior to behavioral assessment, animals were handled daily for 5min to reduce acute anxiety-like responses (44). Testing took place during the light phase and was conducted in accordance with local ethical commission on animal experiment (see supplementary for details). To induce a cognitive deficit in young mice, animals were subjected to a chronic stress protocol (CS). They were placed, twice a day for 1hr, in a 50ml Falcon® tube perforated on each end to let the animal breathe. CS was applied for at least 10 days before testing but was not applied on testing days.

### Ligand preparation and administration

Ligands were diluted in a vehicle solution containing 85% distilled H_2_O, 14% propylene glycol and 1% Tween-80 to be administered intraperitoneally (i.p.) at a volume of 10ml/kg. Working solutions were prepared at 1, 5 or 10mg/kg and adjusted to body weight before injection. DZP was used as a non-selective GABAA-R PAM with known anxiolytic activity, which does not exert known pro-cognitive effects. DZP was administered i.p. at 1.5mg/kg based on previous studies (45).

For sub-chronic administration in the drinking water for 10 consecutive days, ligands were diluted in tap-water, stirred overnight at room temperature and provided in glass bottles to tailor for a 30mg/kg daily dose. Bottles were changed every other day to provide freshly-prepared solutions.

### Pharmacokinetic characterization

Metabolism studies were performed in human and mouse liver microsomes as described in (46). Briefly, the test ligands were incubated at 10µM with active or heat-inactivated human or mouse liver microsomes and appropriate cofactors. Aliquots were removed at different time points and assayed using a liquid chromatography/tandem mass spectrometry (LC-MS/MS) analytical method. To obtain pharmacokinetic profiles, mice were treated i.p. at 10mg/kg and sacrificed at different time points (5-to-720min post-injection) for brain and plasma quantification of ligands by ultraperformance LC–MS/MS (47).

### In vitro hydrolytic plasma stability studies

Each ligand stability was tested *in vitro* at 37 °C, utilizing blank mouse plasma spiked with the respective ligand and internal standard, as described in (39).

### Plasma protein and brain tissue binding studies

The rapid equilibrium dialysis assay used to determine the free fraction of the ligands in mouse plasma and brain tissue was as described in (47). Free concentrations in the brain were calculated by multiplying the obtained total brain concentrations with the appropriate free fractions determined by rapid equilibrium dialysis.

### Behavioral assessment

The ligands and the non-selective PAM DZP were tested in behavioral tests assessing anxiety-like behavior (elevated plus maze, **EPM**), antidepressant predictability (forced swim test, **FST**), locomotor activity (open-field), motor coordination (rotarod), and spatial-working memory (Y-maze, **YM**). See details in the **Supplementary methods** and **Supplementary Figure S2**.

### Statistical analysis

All data are expressed as mean ± SEM. Data obtained from electrophysiology studies are subjected to t-tests comparing the mean to 100% of response to GABA alone. For behavioral experiments, statistical analyses were performed using one-way ANOVA and post-hoc Scheffe’s or Student-Newman-Keuls test when applicable. For the EPM and YM tests, sex was put as a cofactor to investigate potential sex effects on the behavioral outcome. All values obtained after statistical analysis are provided in the **Supplementary Tables S1, S2, S8, S9, S10, and S11**.

## RESULTS

### Novel IBZD amide ligands potentiate α-containing GABAA-Rs

To assess potentiation at the different α-containing recombinant GABAA-Rs (**Figure 1a-d and Supplementary Table S1**), the effects of four new IBZD amide ligands were compared to the response to GABA alone (EC_3-5_). At 100nM, GL-II-74 and GL-II-75 exhibit substantial allosteric modulation at α5-GABAA-Rs, while GL-II-73 and GL-II-76 exhibited lower, but still significant, potentiation (t-test comparison to 100%; t>2.9; p<0.04). At this low concentration, all compounds also potentiate α1, α2 and α3-GABAA-Rs (t>2.9; p<0.04), GL-II-75 already exhibiting high potentiation at this subunits, suggesting shared electrophysiological properties with the referent non-selective PAM DZP (48,49). The profiles at 100nM were confirmed at 1µM concentration, with strong α5-PAM effects (t-test comparison to 100%; t>4.67; p<0.009), and still significant potentiation at α1,α2 and α3 (t>3.17; p<0.03). ANOVA comparing potentiations between all subunits for each compound at each concentration revealed that all compounds but GL-II-75 preferentially potentiate α5-GABAA-Rs (ps<0.04; **Supplementary Table S2**) As expected, α4-, α6- or δ-subunit containing GABAA-Rs were not potentiated by the new ligands (**Supplementary Table S1**) suggesting similar subunit-dependent activity as DZP (48,50-52). Similar potentiation levels were obtained at the β1- and β3-containing GABA_A_ receptors (besides a significant difference for GL-II-73 at 1µM concentration (p=0.03), but irrelevant for in vivo application) suggesting no critical impact of the β-subunit on the modulatory effects of the new ligands (**Figure 1e-h** and **Supplementary Table S3**).

**Figure 1.**
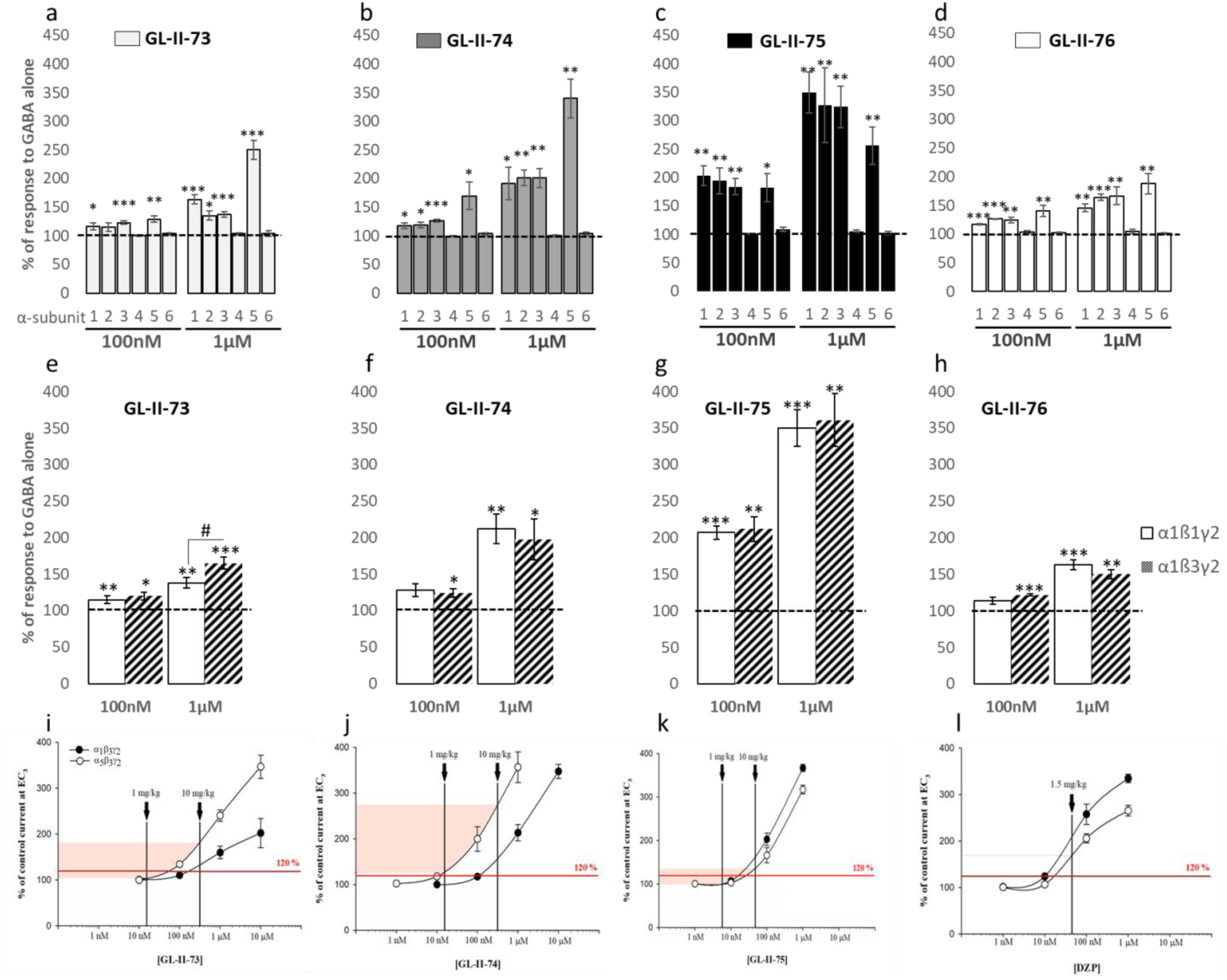
Electrophysiological recordings at GABA_A_ receptors. Modulation properties of 100nM or 1µM of GL-II-73 (**A,E**), GL-II-74 (**B,F**), GL-II-75 (**C,G**) or GL-II-76 (**D,H**) at recombinant α1/2/3/4/5/6β3γ2 (a-d) or α1β1/3γ2 (e-h) receptors. The potentiation of GABA alone (EC_3-5_ GABA) is expressed in percentage. *p< 0.05; **p < 0.01 and ***p < 0.001 compared to 100%, #p<0.05 compared to α1β3γ2. The approximated electrophysiological responses elicited by the estimated free brain concentrations and presented on the concentration-response curves of GL-II-73 (**I**), GL-II-74 (**J**), GL-II-75 (**K**) and DZP (**L**) at rat recombinant α1β3γ2 and α5β3γ2 GABAA receptors measured at GABA EC3 (eliciting 3% of the maximal GABA current in the respective subtype). Brain tissue density of 1.04 g/ml was used to convert brain concentrations from ng/g into ng/ml. The shaded range delineates the interpolated lower and upper limit of potentiation at α5GABA_A_Rs in the dose range used; the vertical lines mark the approximated free brain concentration of the given dose of each ligand. The level of potentiation of 120% was arbitrarily set as borderline for eliciting in vivo behavioral effects.

### Ligands selection and doses validation for animal studies

Liver microsome metabolism assays (**Supplementary Table S4**) indicated that GL-II-73 and GL-II-74 had the longest half-lives in mouse and human, respectively, while GL-II-76 had a very short half-life in mice, precluding its use in vivo, hence it was excluded from further analysis.

Considering the potential implication of the α5-subunit on cognitive functions, and the role of α1-subunit in sedation, we aimed at finding a dose that would potentiate α5-GABAA-Rs preferentially to α1-GABAA-Rs (26). The concentration-response curves for the ligands (**Figure 1i-k** and **Supplementary Table S5**) suggest that at 1mg/kg, expected potentiation would be below 120% for GL-II-73 and GL-II-75 at both α1- and α5-GABAA-Rs, while GL-II-74 would have significant potentiation at α5-GABAA-Rs. At 10mg/kg, potentiation would significantly increase for both subunits, with higher potentiation at α5-than α1-GABAA-Rs for GL-II-73 and GL-II-74. Conversely, GL-II-75 exhibits higher potentiation at α1-than at α5-GABAA-Rs at 10mg/kg. Higher potentiation at α1-than α5-GABAA-Rs was also observed with the non-selective referent DZP (**Figure 1l**).

Potentiation of GABA-induced chloride current does not necessarily correlate with binding affinity. Indeed, binding assays at α1/2/3/5-GABAA-Rs showed that all ligands are affinity-selective to the α5-GABAA-R (Ki values being 6-15x more potent at α5-GABAA-R than other α-GABAARs, **Supplementary Table S6**). Notably, GL-II-73 showed the lowest affinities, with Ki values in the µM range, compared to the other ligands. In contrast to all four novel ligands, DZP showed high and similar affinities at all α-GABAA-Rs.

Together, these results establish GL-II-73, GL-II-74 and GL-II-75 as ligands with significant PAM efficacies at α5-GABAA-Rs, and low, moderate and higher efficacies at α1/2/3-GABAA-Rs, respectively, and also as α5-GABAA-R-binding preferring ligands, possessing relatively high affinities, with the notable exception of the low µM range for GL-II-73.

### Pharmacokinetic, plasma stability and free fraction studies

Pharmacokinetic profiles of the test ligands and DZP were established in mice after 10mg/kg i.p. administration (**Figure 2a-d**). Plasma and brain free fractions were 20.39% and 12.14% for GL-II-73, 6.34% and 4.49% for GL-II-74, 5.08% and 1.34% for GL-II-75, and 16.25% and 6.38% for DZP, respectively. Elimination half-lives (T_1/2_) of all three ligands and DZP suggest that DZP and GL-II-74 are the most stable in mouse plasma, while GL-II-73 is most stable in the mouse brain.

**Figure 2.**
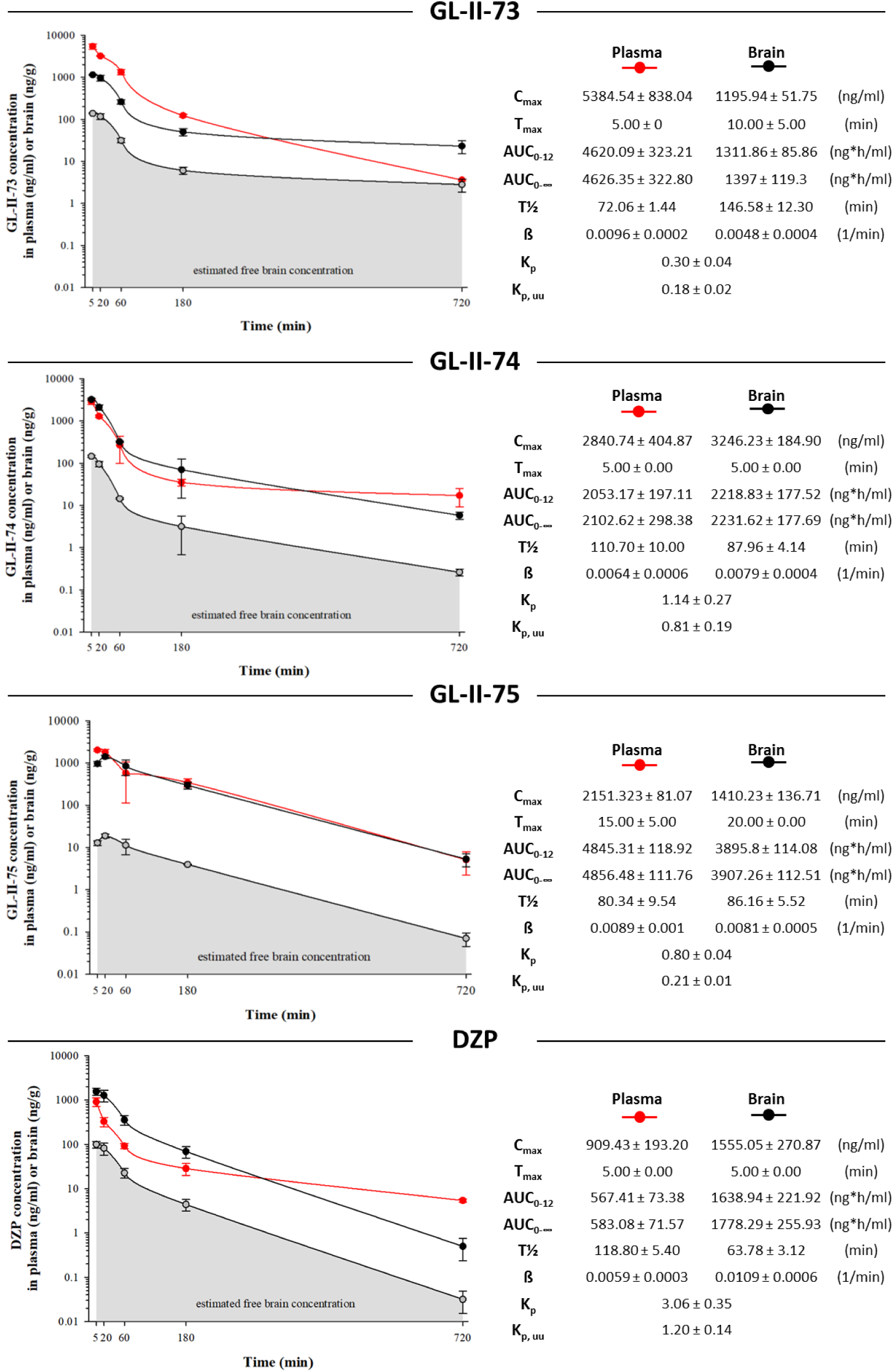
Plasma and brain concentration–time profiles of GL-II-73, GL-II-74, GL-II-75 and DZP. Plasma and brain concentration–time profiles of GL-II-73, GL-II-74, GL-II-75 and DZP after intraperitoneal administration of the 10 mg/kg dose in male C57BL/6 mice (n=3 per time point). C_max_ = maximum concentration in plasma or brain; T_max_ = time of maximum concentration in plasma or brain; AUC_0-720_ = area under the plasma or brain concentration–time curve from 0 to 720 min; AUC_0-∞_ = area under the plasma or brain concentration–time curve from 0 to extrapolated infinite time; t_1/2_ = elimination half-life from plasma or brain; β = elimination constant rate from plasma or brain; K_p_ = brain-to-plasma partition coefficient (K_p_ = AUC_0-∞_, brain/AUC_0-∞_, plasma); K_p,uu,brain_ = ratio of unbound brain to unbound plasma drug concentrations (K_p,uu,brain_ = Kp × unbound fraction in brain/unbound fraction in plasma). All values are represented as mean ± standard error of the mean.

Maximum brain concentration (C_max_) was the highest with GL-II-74, suggesting a relatively optimized capacity for brain targeting. Brain-to-plasma partition coefficient (K_p_) values showed that GL-II-74 displayed excellent brain permeability, although less efficient than DZP (Kp=1.14 vs 3.06). This is further supported by the ratio of unbound brain to unbound plasma ligand concentrations values (K_p,uu_), a measure of net transport across the blood-brain barrier, which better quantifies the brain penetration efficiency (53). This parameter demonstrates that GL-II-73 may be a substrate for efflux transport mechanisms at the blood-brain barrier, compared to GL-II-74 or GL-II-75. Although DZP displayed excellent brain permeability (as measured by K_p_ and K_p,uu_), GL-II-74 and GL-II-75 reached notably higher AUC brain values (from 0 to 720 min or infinite time post-dosing) than DZP. The brain AUC value for GL-II-73 was somewhat lower but comparable to DZP.

All three ligands displayed acceptable brain penetration and an excellent *in vitro* metabolic stability; after 4h of incubation in mouse plasma, the fraction of remaining intact ligand was 98.88%, 85.83% and 78.52% for GL-II-73, GL-II-74 and GL-II-75, respectively, whereas the intact fraction of DZP was only 76.36%. Thus, the novel ligands have at least as favorable pharmacokinetic properties as DZP.

The concentrations of all ligands in mouse brain 30 min after single, or 1h after the last of three i.p. injections (administered 24h, 20h and 1h prior to determination of concentration) at 1, 5 and 10mg/kg (**Supplementary Figure S2** and **Supplementary Table S7**) demonstrated that attainable concentrations were governed by the dose and dosage regimen. In view of the potentiation at α1- and α5–GABAA-Rs (**Figure 1**) and pharmacokinetic properties (**Figure 2**), doses of 1, 5 and 10mg/kg doses were chosen for behavioral testing.

### Estimated in vivo receptor occupancy

The relative fraction of receptors occupied at the given free concentration can be approximated from in vitro binding and PK data (Supplementary **Table S7**). At 10mg/kg, GL-II-73 displayed very low predicted receptor occupancy, including at the α5-GABAA-R (5.95%). GL-II-75 displayed low values at α1/2/3-GABAA-Rs (<9.31%), and 38.42% predicted occupancy at α5-GABAA-Rs. GL-II-74 displayed values in the 17-24% range for α1/2/3-GABAA-Rs and 75.51% for α5-GABAA-Rs. In contrast, DZP displayed a much greater overall occupancy at all four α5-GABAA-R (70% and above), even at the 1.5mg/kg low dose. Despite these variable predicted receptor occupancy levels, we used the electrophysiological responses at α1- and α5-GABAA-Rs as guide for in vivo studies.

### Effects of novel IBZD amide ligands with α5-PAM activity on locomotor activity

The effects of GL-II-73, GL-II-74 and GL-II-75, dosed i.p. at 10mg/kg were assessed on the distance traveled in 5-min bins in the locomotor activity test in mice (**Supplementary Figure S3**). A two-way repeated-measure ANOVA, followed by post-hoc comparison revealed that GL-II-74 induced a hyperlocomotion response in time intervals 0-5 min and 50-55 min post-injection, while GL-II-75 elicited a similar stimulant-like effect in the first 5 min, but a consistent hypolocomotion in the 15-40 min period. DZP dosed at 1.5 mg/kg i.p. induced a hypolocomotor response in intervals 10-35 min and 40-55 min (**Supplementary Figure S4)**, confirming its known sedative effect on spontaneous locomotor activity. At contrary, GL-II-73 had no effect on locomotor activity.

### The novel IBDZ amide ligands exhibit anxiolytic and antidepressant-predictive properties

In the EPM test, all ligands induced a trend or a significant increased percentage of time spent in the open-arms of the EPM (ANOVAs; Fs>3.7; p<0.07) Post-hoc analysis identified a significant increase in time after i.p. administration at 10mg/kg (p<0.03) for all compounds(**Figure 3a-d** and **Supplementary Table S9**). 1.5mg/kg DZP significantly increased the time spent in the open-arms (p=0.02), suggesting anxiolytic properties of all ligands, including DZP.

**Figure 3.**
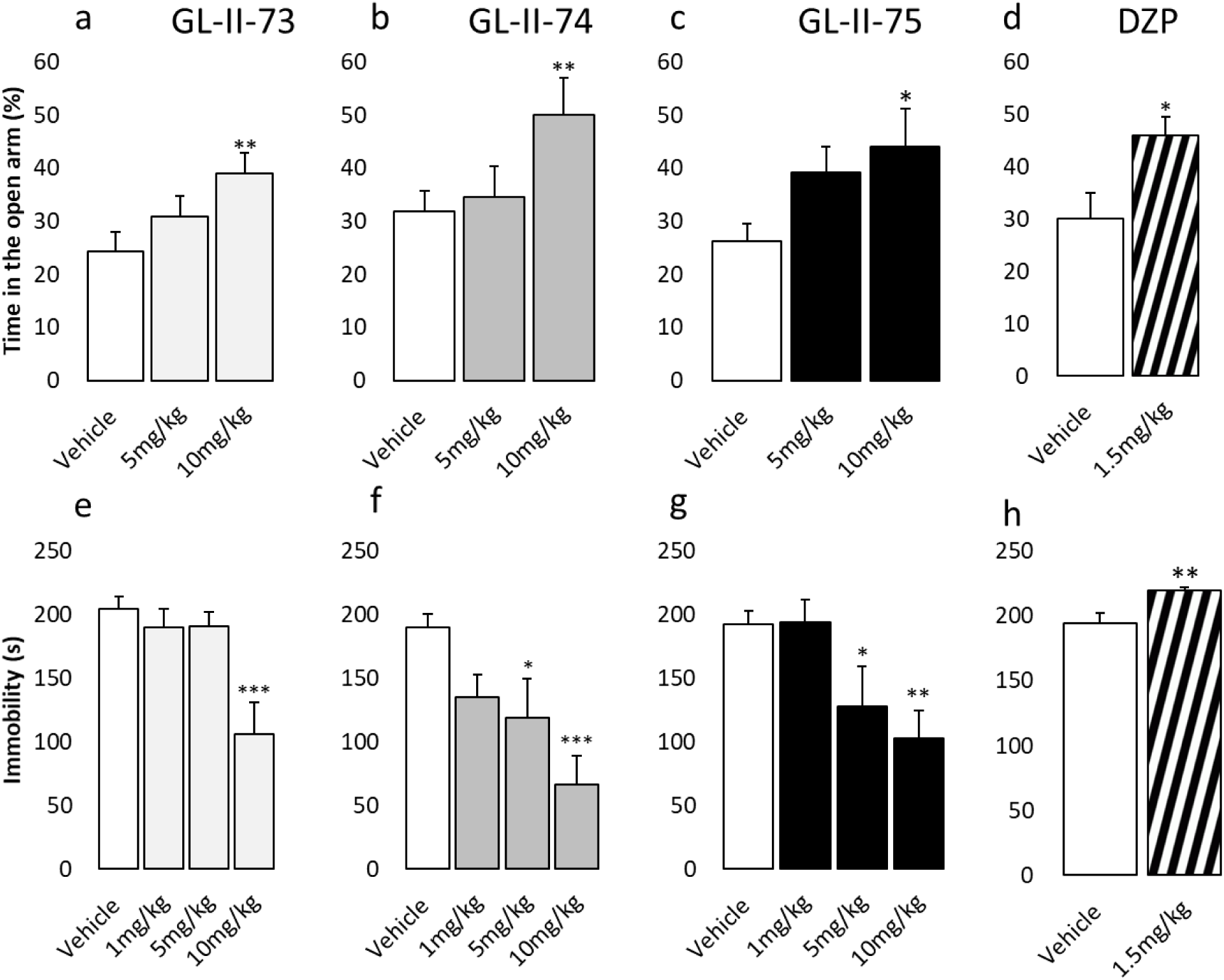
Anxiolytic and antidepressant properties of the α-PAM compared to diazepam. Potential anxiolytic action of the new IBZD amide ligands GL-II-73 (**A**), GL-II-74 (**B**) and GL-II-75(**C**) at 5 or 10 mg/kg and the referent non-selective PAM diazepam (DZP: 1.5mg/kg; **D**) has been assessed in mice (50% females) using the elevated plus maze. Animals received vehicle or the ligand GL-II-73 (n(0)=13, n(5)=13 and n(10)=14), GL-II-74 (n(0)=14, n(5)=13 and n(10)=14), GL-II-75 (n(0)=13, n(5)=14 and n(10)=13) or DZP (n(0)=11 and n(1.5)=10), 30 minutes before testing. A significant increase in the time spent in the open arms was used as an index of potential anxiolytic action. Potential antidepressant properties of the ligands were assessed in male mice using the forced-swim test (**E-H**). Sex as a cofactor was not significant (p-values≥0.17). Mice were placed in an inescapable transparent tank filled with water (25cm, 26±1^°^C) for a period of 6min. Immobility is defined as the minimum amount of movement to stay afloat, between the second and the sixth minute of testing. Mice were injected following serial i.p. administrations at 1, 5 or 10 mg/kg for the new ligands (GL-II-73: n(0)=8, n(1)=8; n(5)=6 and n(10)=8; GL-II-74: n(0)=8, n(1)= 8, n(5)=8 and n(10)=8; GL-II-75: n(0)=8, n(1)=8, n(5)=8 and n(10)=9) or 1.5mg/kg for the DZP (n(0)=12 and n(1.5)=12). Significant decreased immobility characterized the potential antidepressant-like efficacy of the ligand. **p < 0.01 and ***p < 0.001 compared to Vehicle, ###p < 0.001 compared to GL-II-73. All values are represented as mean ± standard error of the mean.

In the FST, GL-II-73, GL-II-74 and GL-II-75 induced a significant decrease in time spent immobile (ANOVA F>5.4; p<0.004) compared to the vehicle-injected group after i.p. administration at 10mg/kg (p<0.003; **Figure 3e-h**). The dose of 5mg/kg also reduced the time spent immobile in animals receiving GL-II-74 and GL-II-75 (p<0.03). In contrast, DZP induced significant increases in time spent immobile (p=0.004), potentially due to locomotor side-effect (**Supplementary Figures S5**), thus precluding any conclusion as to putative depressant-like effect.

### The novel IBZD amide ligands reverse stress-induced working memory deficits

ANOVAs reveled differences in alternation rate after CS exposure and injection of the ligands (F>7; p<0.0004). Young adult mice were exposed to chronic stress (CS) to induce working memory impairments in a YM spatial spontaneous alternation task. CS exposure decreased the alternation rate in animals receiving only vehicle (p<0.002; **Figures 4A-D and Supplementary Table S9**). Administration of GL-II-73 at 10mg/kg restored alternation rate in stressed animals to the same level as non-stressed animals, and significantly different from the stressed mice receiving vehicle (p=0.01). 1mg/kg and 5mg/kg doses were inefficient. In contrast, stressed animals injected with GL-II-74 displayed lower alternation rate than non-stressed animals, suggesting a lack of pro-cognitive effect, regardless of the ligand dose (p>0.97). GL-II-75 administration restored alternation rate to non-stressed levels at 5 and 10mg/kg (p<0.045). As expected, the administration of 1.5mg/kg of DZP did not reverse the cognitive deficits induced by CS (p=0.94).

**Figure 4.**
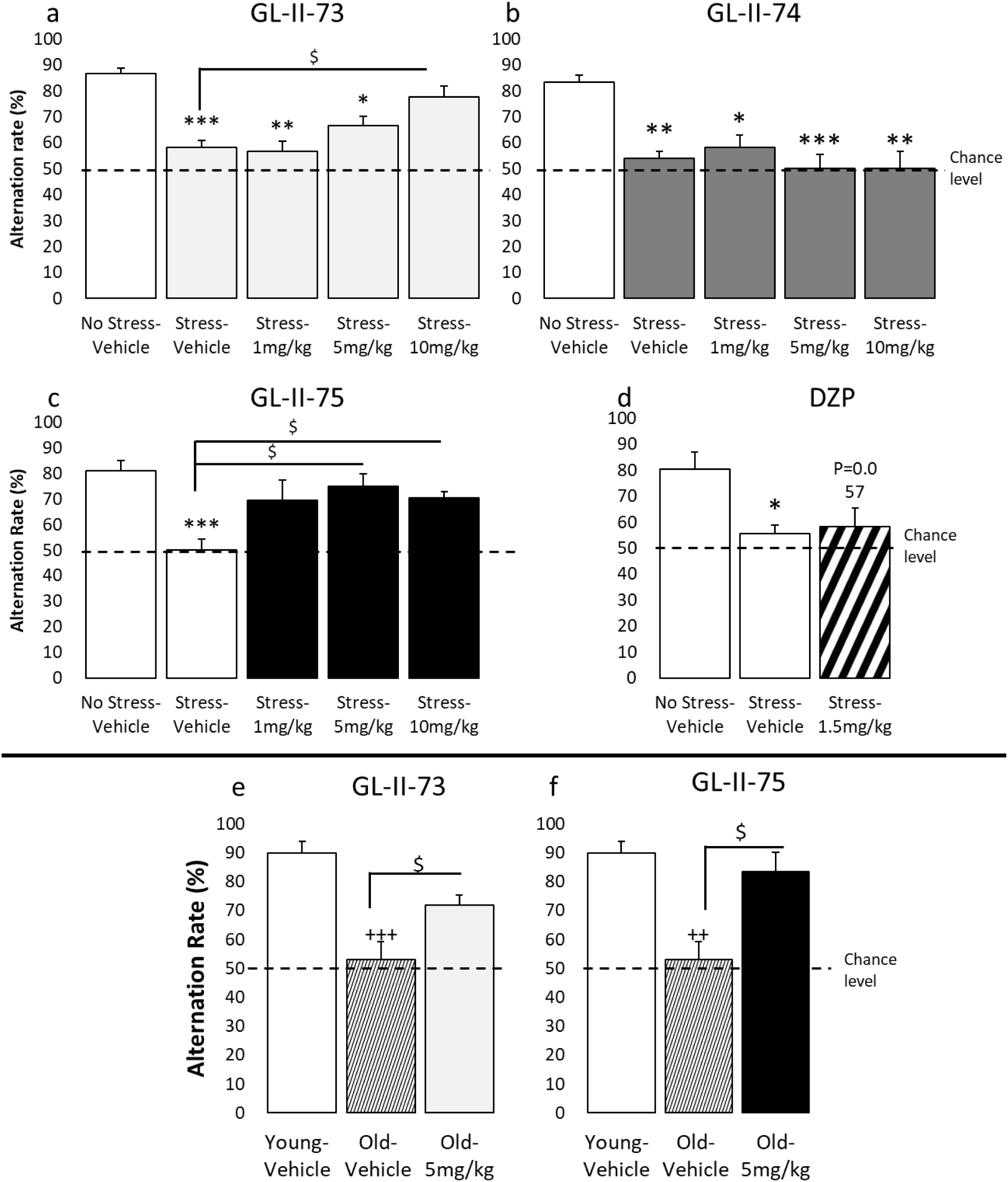
Pro-cognitive action of GL-II-73 and GL-II-75 on stress-induced and age-related working memory impairment. Cognitive effects were assessed in a spontaneous alternation task using a 90sec inter-trial interval (ITI). Prior to experiment with young mice, a cognitive deficit was induced by exposing the animals to daily chronic restraint stress, 1hour twice a day for 1 week. Young mice (50% females) received vehicle or 1, 5 or 10mg/kg of GL-II-73 (**A**; n(0-NS)=10, n(0-S)=10, n(1)=5, n(5)=10, n(10)=12). The same protocol was used for GL-II-74 (**B**; n(0-NS)=8, n(0-S)=8, n(1)=4, n(5)=10, n(10)=4)), GL-II-75 (**C**; n(0-NS)=8, n(0-S)=8, n(1)=6, n(5)=4, n(10)=9) and DZP (**D**;n(0-NS)=6, n(0-S)=6, n(1,5)=6). Animals were injected i.p. with vehicle, α5-PAMs or DZP, 30min prior testing. For old animals (**E-F**), the same protocol was applied with the ITI shortened to 60sec. Old male mice received Vehicle or GL-II-73 (**E**; n(0-Young)=5, n(0-Old)=5, n(5)=6) or GL-II-75 (**F**; n(0-Young)=5, n(0-Old)=5, n(5)=4) and were compared to young and old mice treated with Vehicle. For all experiments described here, sex as a cofactor was not significant (p-value≥0.49). Results are presented as mean of the percentage of alternation ± SEM: effect of the stress: *p<0.05; **p<0.01; ***p<0.001 compared to “No stress-vehicle”; Effect of age: ++p<0.01 or +++p<0.001 compared to “Young-Vehicle”; Effect of the ligand $p<0.05, $$p<0.01 compared to “CRS-Vehicle” or “Old Vehicle”.

We also tested the ligands in non-stressed animals to assess putative effects at baseline (**Supplementary Figure S6** and **Supplementary Table S9**). GL-II-73 and GL-II-75 had no effect on alternation rate (ANOVAs F<2.4, p>0.3), whereas 10mg/kg of GL-II-74 and 1.5mg/kg DZP reduced alternation rates (ANOVAs F>8.8; p<0.005), suggesting deleterious effects on working/spatial memory.

### GL-II-73 and GL-II-75 reverse age-related working memory deficits

The pro-cognitive efficacy of GL-II-73 and GL-II-75 was assessed in old male mice. ANOVAs reveled significant differences between young, old and old treated (F<0.0015). 18-month old mice displayed alternation rates in the Y-maze at chance level, suggesting cognitive impairment (p<0.002 compared to young mice). A single 5mg/kg administration (i.p) of GL-II-73 or GL-II-75 significantly reversed spatial-working memory deficits of old mice to levels indistinguishable from young controls (p<0.03; **Figure 4E-F and Supplementary Table S10**).

### GL-II-73, but not GL-II-75, maintained pro-cognitive activity after sub-chronic administration in young and old mice

The two ligands that exhibited pro-cognitive efficacy after a single acute i.p. injection (GL-II-73 and GL-II-75) were tested via sub-chronic administration in drinking water (**Figure 5** and **Supplementary Table S11).** In the adult stress-induced cognitive deficit model, ANOVA analysis revealed significant differences between groups (F(2,14)=28.1; p=0.0001) only with GL-II-73 injection, characterized by increased alternation rate after sub-chronic administration (P=0.005 compared to stressed mice). A similar result was obtained in the age-induced cognitive deficit model, where GL-II-73 (p=0.0001) but not GL-II-75 (p=0.46) increased alternation rate after sub-chronic administration.

**Figure 5.**
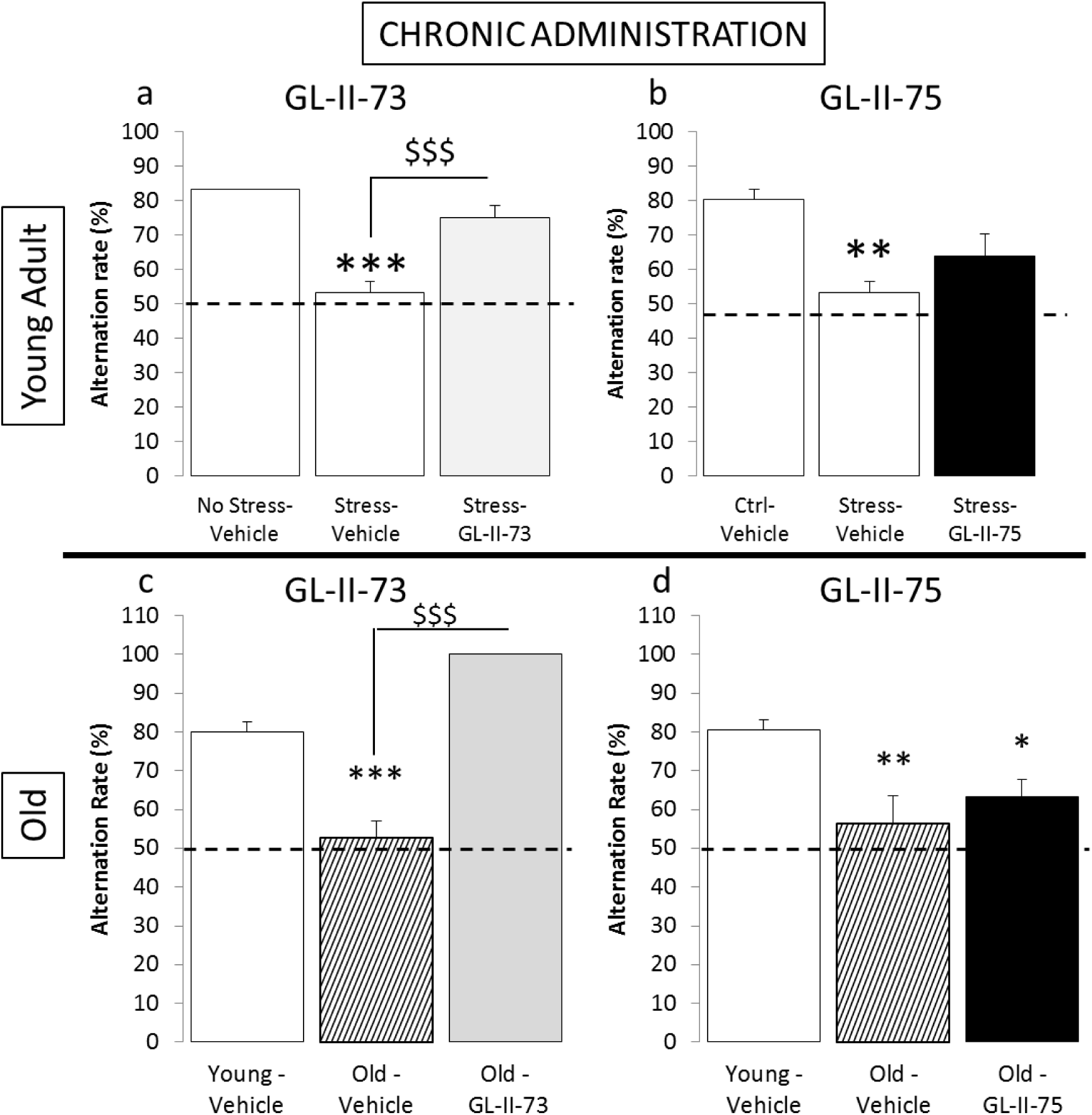
Pro-cognitive action of sub-chronically administered GL-II-73 and GL-II-75 on stress-induced and age-related working memory impairment. Pro-cognitive properties of GL-II-73 (**A, C**) and GL-II-75 (**B, D**) were assessed in young (**A-B**) and old (**C-D**) male mice after sub-chronic administration in the drinking water for 10 days. Cognitive abilities were assessed in a spontaneous alternation task. Alternation rate was calculated as the percentage of correct alternations in function of the maximum alternation possible (i.e. 6). Prior to experiment, a cognitive deficit was induced by exposing the young animals to chronic stress for 1 week. Young animals received sub-chronically GL-II-73 (n(0-NS)=6, n(0-S)=5, n(30)=6) or GL-II-75 (n(0-NS)=5, n(0-S)=5, n(5)=6) dosed at 30mg/kg. Old mice received sub-chronically GL-II-73 (n(0-Young)=5, n(0-Old)=6, n(30)=4) or GL-II-75 (n(0-Young)=6, n(0-Old)=5, n(30)=5) dosed at 30mg/kg in the drinking water. Results are presented as mean of the percentage of alternation ± SEM: effect of the stress: *p<0.05; **p<0.01;***p<0.001 compared to “No stress-vehicle”; Effect of age: ++p<0.01 or +++p<0.001 compared to “Young-Vehicle”; Effect of the ligand $p<0.05, $$p<0.01 compared to “CS-Vehicle” or “Old Vehicle”.

## DISCUSSION

The present study identified for the first time the combined antidepressant, anxiolytic and pro-cognitive properties of newly designed IBDZ amide ligands with α5-PAM activity. This work is based on the hypotheses that deficits in signaling through these receptors contribute to mood and cognitive symptoms in depression and during aging, and that selective enhancement of activity at α5-GABAA-Rs may have therapeutic potential, compared to non-selective BZs that lack such activities. We generated four ligands; metabolic stability of one of them (GL-II-76) was unfavorable, potentially due to the pyrrolidine residue in the amide moiety, and it was not tested further. Binding and electrophysiological assessment indicated that GL-II-73 and GL-II-74 acted as PAMs with priority affinity and efficacy at α5-GABAA-Rs, whereas GL-II-75 potentiated GABA-gated chloride current to a greater extent at α1-, α2- and α3-GABAA-Rs compared to α5-GABAA-R, suggesting properties closer to DZP (48), although with much lower affinity at α-GABAA-Rs than DZP. The lack of potentiation at GABAA-Rs containing the α4, α6 or δ subunits (54) supports the notion that all ligands bind to the DZP-specific BZ-sensitive site of GABAA-Rs.

Brain and plasma pharmacokinetic studies showed that the ligands are brain-penetrant, with penetration indices (K_p_, K_p,uu_) ranked in the following decreasing order: DZP > GL-II-74 > GL-II-75 > GL-II-73. Accordingly, all ligands were tested for behavioral activity in vivo. The three newly-designed ligands as well as the non-selective DZP (55) exerted an anxiolytic effect, demonstrating conserved anxiolytic effects despite variable efficacy profiles, probably attributable to a sufficient degree of modulatory activity at α2-GABAA-Rs (27). In the forced-swim test, all three new ligands displayed antidepressant-like properties, at doses that induced no or low locomotor effects. GL-II-74 and GL-II-75 demonstrated altered locomotor activity and diminished coordination at the higher doses (10mg/kg), suggesting motor-impairing side-effects, probably due to their higher potentiation at α1-GABAA-Rs (56,57). On the other hand, the affinity of GL-II-73 for α1-GABAA-Rs is exceptionally low, which profoundly decreases the propensity of this ligand to elicit any motor impairment. In contrast, DZP increased the time spent immobile, confirming a lack of antidepressant effect, or even an induction of depressant-like state, although the latter was probably confounded by motor-impairing effects (58).

We tested the potential of novel ligands at reversing spatial working memory deficits induced by chronic stress exposure(14,59) or related to normal aging (60,61). GL-II-73 and GL-II-75 exhibited significant pro-cognitive efficacy in the stress-induced cognitive deficit model, whereas GL-II-74 displayed no pro-cognitive effects. Notably, the pro-cognitive effects of GL-II-73 and GL-II-75 were confirmed in aged mice, as old animals performed at levels observed in young animals after a single injection. The pro-cognitive effect of GL-II-73, but not GL-II-75, was maintained after sub-chronic administration in both the stress-induced and aging models. It remains to be tested whether this difference reflected the shorter brain half-life or other kinetic parameters of GL-II-75, or a potential desensitization effect. Consistent with previous findings, DZP had no pro-cognitive effects (62). Together, these results demonstrate for the first time the potential for anxiolytic and combined antidepressant and pro-cognitive properties of newly designed IBDZ amide ligands with efficacy at α5-GABAA-Rs.

It was recently shown that co-localization of the α1-, α5- and γ2-subunit in the hippocampus is necessary for successful spatial learning (26). From this postulate, the potentiation at both α5- and α1-GABAA-Rs could explain pro-cognitive properties of GL-II-73 and GL-II-75 after acute/sub-chronic administration in our working memory task, however it does not explain the lack of pro-cognitive effects of GL-II-74 and of BZs in general. Moreover, it was shown in experiments combining DZP with α5- or α1-GABAA-R antagonists that DZP incapacitation action in the Morris water maze, a reference spatial learning and memory test (63), are mediated by α1-GABAAR activation and that blockade of α5-GABAAR activity further worsen this incapacitation.

Could all these apparently disparate findings be reconciled in a consistent way in light of other parameters tested or estimated in this study? We suggest that the relative fraction of receptors occupied may be a relevant factor (**Table 1**). Despite significant efficacy and robust behavioral effects, GL-II-73 displayed unprecedented low overall receptor occupancies, compared to GL-II-74, GL-II-75 and DZP. This suggests that a low fractional occupancy at α1-GABAA-Rs (<10%, and preferentially <1%), accompanied by a moderate level of positive modulation at both α1-and α5-GABAA-Rs, gives rise to an improved side-effect profile, together unmasking antidepressant or pro-cognitive potential of α5-GABAA-Rs. Hence, high fractional occupation may preclude any antidepressant or pro-cognitive properties, consistent with the prodepressant-like(55) and amnesic effect of DZP (64) suggested to be mediated by α1-GABAA-Rs (65,66). The role of the α1- and α5-subunit seems to follow a biphasic pattern, where low activation facilitates mood and cognitive processes whereas high and sustained activation impairs these functions (67).

Negative allosteric modulators (NAMs) at GABAA-Rs can also exert antidepressant (68) and pro-cognitive (69,70) activity in tests such as passive avoidance learning, recognition learning (71) and spatial memory (72). Moreover, patients with cognitive impairment in Down syndrome were unsuccessfully clinically-trialed with the α5-selective NAM basmisanil, which is currently being tested for cognitive impairment associated with schizophrenia (73). However, reducing the function of α5-GABAA-Rs is predicted to worsen the pathology associated with reduced SOMATOSTATIN cell function and increased neuronal excitability, hence being potentially associated with risk for long-term detrimental effects, and Alzheimer’s disease development (74). However, the putative pro-cognitive properties of both α5-NAMs and α5-PAMs are not mutually incompatible. α5-NAM could be acutely efficient in certain cognitive tasks such as spatial reference (70) where disinhibition of pyramidal neurons may facilitate the acquisition of a mental spatial map. In contrast, an α5-PAM could be efficient in cognitive tasks (such as working memory) where increased inhibition of neuronal activity may reduce noise and interferences, and increase signal-to-noise ratio for incoming stimuli (6,75), together strengthening the salient inputs that need to be kept for high-level performances (76). This hypothesis on the roles of NAMs and PAMs will require further validation.

Given the increased rates of cognitive and mood impairments with psychiatric diseases and aging, it is imperative to develop new therapeutic strategies based on emerging knowledge of primary pathologies of the diseases. Here, we designed, developed, tested and validated preclinical efficacies of novel IBZD amide ligands acting either preferentially (GL-II-73 and GL-II-74) or non-selectively (GL-II-75) as α5-PAM at GABAA-R, hence bypassing the putative SOMATOSTATIN-cell GABA deficit by acting on post-synaptic sites. The fact that these ligands are synthetized from the hybrid DZP/flumazenil privilege structure with very low toxicity profiles and exhibit combined antidepressant and pro-cognitive therapeutic potential (for GL-II-73 and GL-II-75), suggest they may have clinical potential through simultaneously targeting mood and cognition, therefore reducing cognitive deficits associated with depression and other psychiatric disorders, as well as normal aging.

## ACKNOWLEDGEMENTS

We acknowledge the University of Wisconsin-Milwaukee’s Shimadzu Laboratory for Advanced and Applied Analytical Chemistry, the NIH (R01MH096463: R01NS076517), the Ministry of Education, Science and Technological Development, R. Serbia (Grant No. 175076), and the Brain & Behavior Resrach Foundation, awarding NARSAD grants (ES, MB) for generous financial support. We thank the Milwaukee Institute of Drug Design.

